# Biosensor libraries harness large classes of binding domains for allosteric transcription regulators

**DOI:** 10.1101/193029

**Authors:** Javier F. Juárez, Begoña Lecube-Azpeitia, Stuart L. Brown, George M. Church

**Affiliations:** Department of Genetics, Harvard Medical School, Boston, Massachusetts 02115, USA.

## Abstract

Bacteria’s ability to specifically sense small molecules in their environment and trigger metabolic responses in accordance is an invaluable biotechnological resource. While many transcription factors (TFs) mediating these processes have been studied, only a handful has been leveraged for molecular biology applications. To expand this panel of biotechnologically important sensors here we present a strategy for the construction and testing of chimeric TF libraries, based on the fusion of highly soluble periplasmic binding proteins (PBPs) with DNA-binding domains (DBDs). We validated this strategy by constructing and functionally testing two unique sense-and-response regulators for benzoate, an environmentally and industrially relevant metabolite. This work will enable the development of tailored biosensors for synthetic regulatory circuits.

## INTRODUCTION

Synthetic Biology has been a rapidly evolving discipline during the last decades ^1^, propelled by the advances in the synthesis and assembly of ever longer and more complex DNA sequences ^2,3^. This progress in synthesis has been accompanied since the turn of the century by the discovery, characterization and adaptation of novel systems for regulation of gene expression, such as riboswitches ^4–6^, TALENs ^7,8^ and RNA-guided nucleases as CRISPRi-dCas9 ^8,9^. However, the workhorses of the gene regulation world are still monogenic transcription factors (TFs) that regulate genes upon binding a small soluble molecule known as the inducer. Bacterial transcriptional repressors such as LacI ^10^ and TetR ^11^ belong to this group of regulators and have represented the preferred TF choice for decades. Robust TFs such as LacI and TetR have been used to pair an inducer and common reporters (e.g. antibiotic resistance, GFP and LacZ) by controlling the promoters that drive their expression. Compared to the two-component signal transduction systems, this arrangement of sensor and effector in one molecule is simpler and more effective ^12^, making monogenic TF ideal ^13^ for whole-cell biosensor applications ^14^. While two-component systems have the ability to detect molecules that cannot be imported into the cytoplasm, their use as biosensors is limited by the risk of cross-talk between them ^15^. In this work, we focus exclusively on the development of monogenic intracellular sensors to avoid the undesired traits of two component systems, such as their expression as membrane proteins and the risk of activation by unspecific phosphorylations.

Despite their advantages, there are only a small number of available monogenic TFs ^16^. Rational design of synthetic monogenic TFs that respond to small molecules of choice has been a long-term aspiration of synthetic biologists and would be an extremely powerful tool for biotechnological applications ^13,17^. In this work, we present a high-throughput pipeline for *in vitro* construction and *in vivo* testing of tailor-made transcriptional regulators by massively multiplexed fusion of protein domains and linkers ^18–20^. Despite knowledge of gene modularity behind the origin of TFs for more than three decades ^21^, the holy grail of a general method for functional fusions of gene domains has remained elusive due to the easily broken allostery ^22^. LacI/GalR regulators can remain active when their substrate-binding domains (SBDs) get swapped within members of the family ^23^. Their DNA-binding domains (DBDs) recognize the same operators but the new TF gets induced by the molecule recognized by the fused SBD. Since the DBDs of the LacI/GalR family were originated from periplasmic binding protein (PBP) that recognized sugars ^24^, there has been at least an attempt to create a novel biosensor substituting LacI-SBD by a PBP. SLCPGL is a glucose-responsive TF built by the fusion of *E. coli* GGBP (Galactose/Glucose Binding Protein) to DBD-LacI ^25^. The chimeric TF Q1 is another example of the generation of a new TF by the fusion of a DBD (from BzdR) to a protein phylogenetically related to its SBD (shikimate kinase) ^26^. The scarce presence in the literature of novel TFs generated by fusing DBD to proteins that are not part of regulators emphasizes the challenge of *de novo* generation of new biosensors.

For the systematic construction of fusion TFs we generated libraries that framed the following degrees of freedom: a) 15 DBDs sourced from bacterial transcriptional repressors with a common architecture and known operator sequences; b) 15 SBDs from Periplasmic Binding Proteins (PBP) associated to ABC transporters. *Figure 1* summarizes the system for the generation of novel TFs described in this work. This pipeline has required the development of new gene assembly methods as well as the construction of a collection of tailor-made reporters. The novel TFs presented here are based on the fusion of DBDs to SBDs that were not part of pre-existing regulators, rather independent proteins with the ability to bind small soluble molecules. DBD and SBD were connected through a series of linkers (LNKs). All the TFs resulting from this combination of domains were functionally tested with benzoate, the inducer molecule specifically recognized by a group of SBDs included in the library. Benzoate was chosen for this proof of concept due to its environmental relevance, as it is associated with the degradation of lignin, the second most abundant polymer on earth after cellulose ^27,28^. The reductive fractionation of lignin frees abundant benzoate-related aromatic compounds that can be used as building blocks for multiple applications, such as biofuel production ^29,30^. On the other hand benzoate is an ideal supplement to culture media given the missing ability of *Escherichia coli* to degrade it ^31^ and the fact that its manipulation is rather safe ^32^. BenM is a transcriptional activator that can bind simultaneously benzoate and *cis,cis*-muconate, but its applicability as biosensor is not perfect due to its complex specificity profile ^33,34^. In this context, the creation of a chimeric TF able to sense benzoate specifically is a viable alternative that can open the door for the routine construction of custom-made TFs.

**Figure 1.**
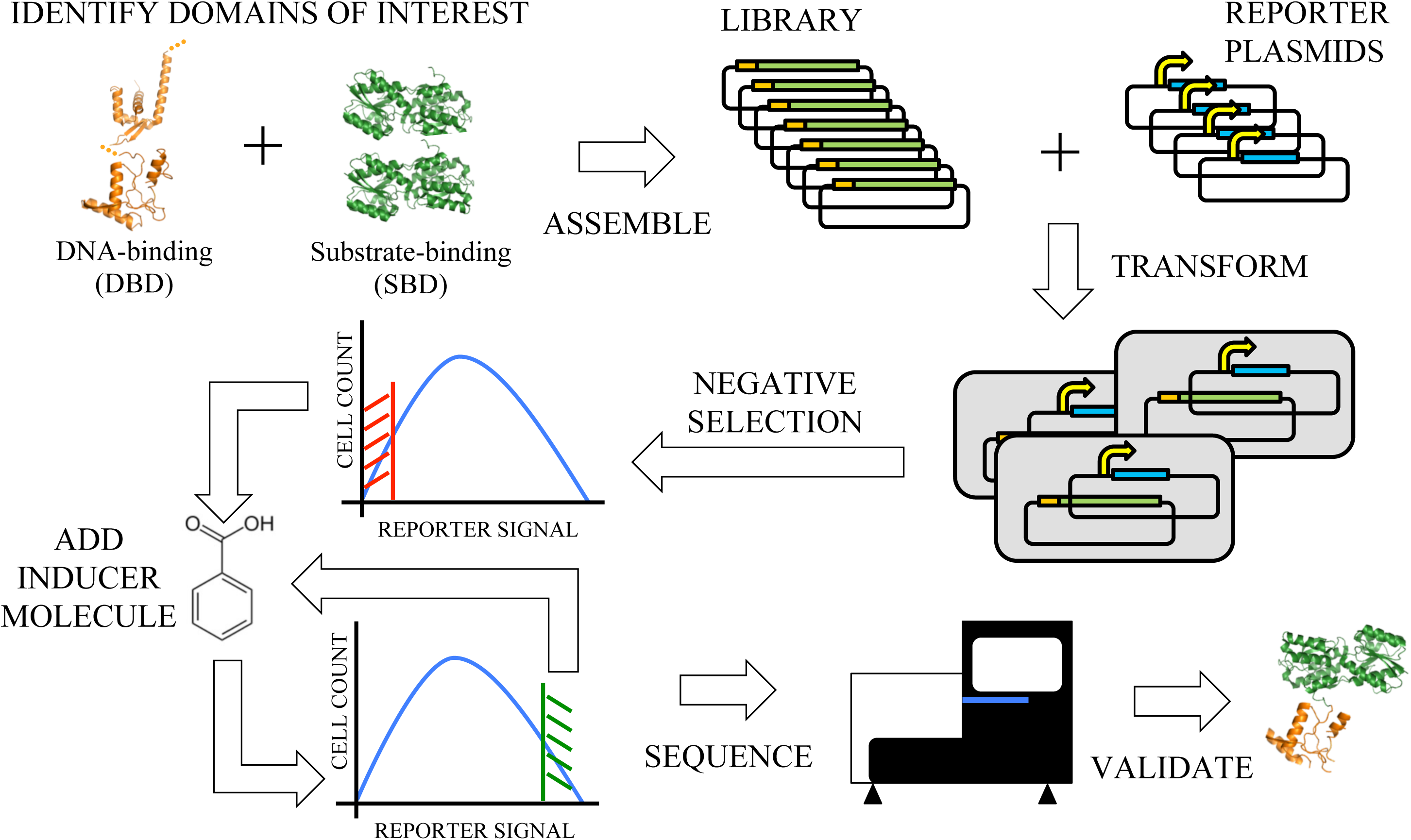
Schematic representation of the pipeline for the generation of novel TFs through gene fusion. Step by step summary outlining the main stages in the generation of chimeric TFs. Starting top-left, the process gets initiated with the selection of the DNA-Binding Domains (DBDs), regions from pre-existing TFs that interact with DNA on specific operator sequences, and the Substrate-Binding Domains (SBDs) that provide the chimera with inducer specificity. Libraries of TFs are assembled combining DBDs and SBDs. In parallel to the assembly of the fusion genes a set of reporter plasmids was developed. Each reporter plasmid contained a modified *Ptac* promoter in which LacI operator boxes had been replaced by operator sequences for one of the DBD included in the collection of chimeras. Both the library of chimeras and the library of reporter plasmids were transformed into *E. coli* host cells. A first negative selection is performed using FACS to enrich the libraries in properly paired DBD/ reporter plasmids: bacteria containing chimeras whose DBD were unable to recognize the operator boxes included in the reporter plasmid could not repress the transcription of a reporter gene encoding GFP. Non-GFP-fluorescent bacteria were recovered and subsequently exposed to the inducer molecule (benzoate). In this case the chimeras that were able to de-repress the reporter promoter allowed the transcription of GFP, facilitating the positive discrimination of inducible chimeras. This positive selection was iterated. As the whole process was performed at library level and we were agnostic about whether we could identify one or multiple functional TFs at the end of the process, the relative abundance of every chimera before and after the selection was compared. To that end both the starting and final libraries were sequenced, and the relative abundance of every chimera was assessed. A set of candidate TFs enriched in the final library was individually assayed *in vivo* to validate its activity. The tridimensional protein structures shown in orange and green correspond to PDB accession codes 1MJL, 3F8C and 2HPH. These structures represent the DNA domains combined in the assembly.

## RESULTS AND DISCUSSION

### Design of the library of transcriptional regulators

We used a novel chimeragenesis method for the construction of custom benzoate-binding TFs. This method is based on the generation of libraries of repressors with a common Nt-DBD-(LNK)-SBD-Ct architecture. All the DBDs were chosen from repressors acting over promoters recognized by the σ^70^ subunit of RNA polymerase and thus carry -10/-35 operator boxes ^35^. The transcriptional repressors inspiring these chimeras share the same domain organization: a Nt region responsible for DNA binding (DBD) is connected to a Ct domain that recognizes a certain compound (SBD), either directly or through linker regions that tend to be less structured than the domains they connect (LNK) ^36^. The native SBD of the TF, which confers the specificity for the inducer molecule is replaced by a protein with the potential to recognize the molecule of interest: benzoate/4-hydroxybenzoate. PBPs were selected to serve as SBDs because they presented multiple advantages: a) high solubility, stability and affinity for their substrates ^37^; b) a detectable conformational change when bound to their ligands ^38^ which is key to change the oligomerization state of the TFs and thus their ability to bind DNA ^39–41^; c) phylogenetic relationship with the SBD of the LacI/GalR family ^24,42^.

Everyone of the resulting chimeric genes carried a DBD, a SBD and, optionally, a LNK between them. Different criteria were set for these three sets of domains, sharing a common goal: maximizing diversity. For DBDs, this objective translated into distinct three-dimensional (3D) configurations, for the LNKs it meant the introduction of different lengths and structural flexibilities. In the case of the SBDs, selection was based on a short list of very similar proteins with demonstrated capability to bind benzoate/4-hydroxybenzoate. This list was expanded to include similar proteins according to amino acid sequence. *Table 1* contains the list of both the transcriptional repressors whose DBD were used for the construction of the chimeric TFs (*Table 1-A*) and the proteins chosen as SBD (*Table 1-B*). The LNK elements incorporated as bridges between DBD and SBD can be found in *Table S1*. Beyond the LNK, another source of diversity in the connection between domains was the inclusion of different variants of every DBD in the library. These variants of a given DBD differed in the last amino acids at the Ct of the domain. The goal was to compensate for the difficulties drawing the borders between domains in the TFs. The shorter version of every DBD was named the “CORE” and the extended variants the “ENDS”. On average there were eight “ENDS” per DBD. On the other hand, two versions of every SBD were included: with and without their signal peptide (SP). A complete description of the selection process and restrictions applied to the final lists of modules for the chimeragenesis can be found in *Supplementary Materials* accompanied by an expanded table including extra information (*Table S1*).

**Table 1.**
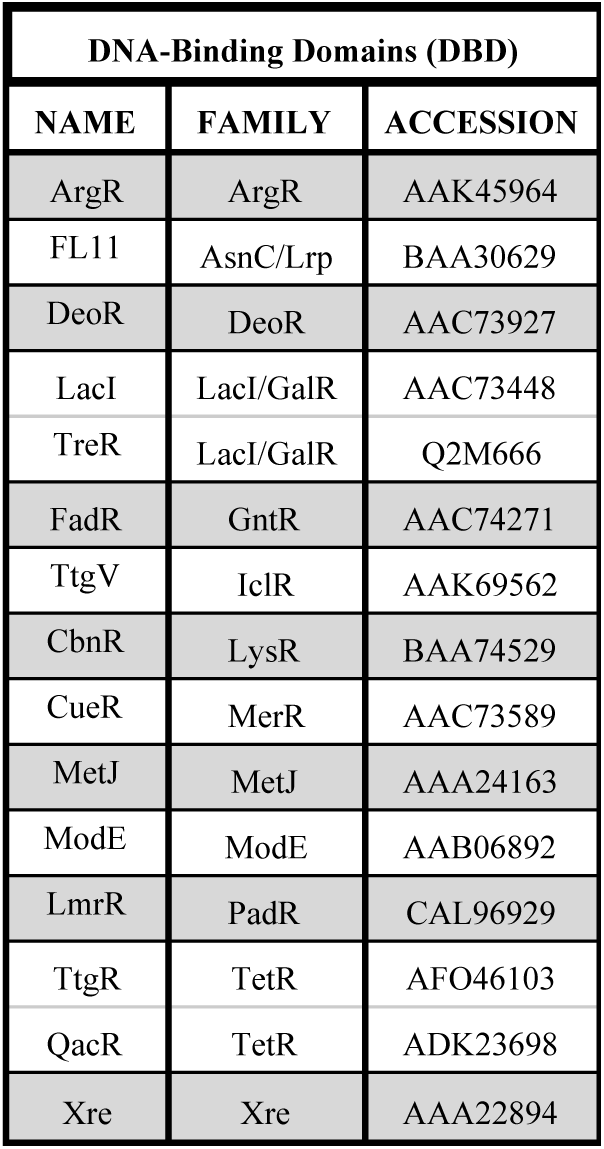
A) DNA-Binding Domains

**Table 1.** B) Substrate-Binding Domains

| Substrate-Binding Domains (SBD) |  |  |  |  |
| --- | --- | --- | --- | --- |
| NAME | ACCESSION | LIGAND | COMMENTS | SIGNAL PEPTIDE (AA) |
| GGBP | AAC75211 | Glucose(*) | Internal Positive Control | 1-23 |
| RPA0668 | CAE26112 | Benzoate | Empirically proven: <sup>84</sup> | 1-31 |
| RPA4029 | CAE29470 | 4-OH-benzoate | Empirically proven: <sup>84</sup> | 1-27 |
| RPA0985 | CAE26428 | 4-OH-benzoate | Empirically proven: <sup>84</sup> | 1-22 |
| ADP71087 | ADP71087 | Benzoate(P) | PID 70% to RPA0668 | 1-31 |
| KAI94709 | KAI94709 | Benzoate(P) | PID 70% to RPA0668 | 1-31 |
| ABE44898 | ABE44898 | Benzoate(P) | PID 53% to RPA0668 | 1-30(E) |
| ABE38823 | ABE38823 | 4-OH-benzoate(P) | PID 89% to RPA4029 | 1-27 |
| EYC50849 | EYC50849 | 4-OH-benzoate(P) | PID 68% to RPA4029 | 1-25 |
| ABD68043 | ABD68043 | 4-OH-benzoate(P) | PID 65% to RPA4029 | 1-24 |
| AHF85493 | AHF85493 | 4-OH-benzoate(P) | PID 65% to RPA0985 | 1-20 |
| CAK09396 | CAK09396 | 4-OH-benzoate(P) | PID 66% to RPA0985 | 1-20 |
| AEK56128 | AEK56128 | 4-OH-benzoate(P) | PID 63% to RPA0985 | 1-29 |
| BzdB1 | AKU11518 | Benzoate(P) | Associated to benzoate catabolism | 1-30 |
| BAE51678 | BAE51678 | Benzoate(P) | Associated to benzoate catabolism | 1-41 |

(\*) Built-in control; (P) Putatively; (E) Estimated, signal peptide identified by similarity to other proteins, not easy prediction based on software described on *Methods*; PID: Percentage of identity (NCBI Protein BLAST)

Finally, an additional domain from LacI was included in the construction of the library with the objective to improve allosteric interactions. The functionality of LacI not only depends on its ability to associate into oligomeric states but also on how long it can maintain the dimerized state ^43^. LacI actually contains at its Ct a so-called oligomerization domain (OD) whose mission is the association of two LacI dimers, creating *de facto* a tetramer that allows it to remain dimerized for longer times, thus increasing its repressing efficiency ^44,45^. The LacI OD was included in the design of the TFs presented in this work so that for every chimera there would be an equivalent carrying an OD translational fusion at Ct. *Supplementary Materials* contain more information on the inclusion of LacI OD as another module of the synthetic TFs.

To identify the different chimeras created in this work we named them by the succession of their elements, from Nt to Ct: DBD-(LNK)-SBD-(OD). Brackets isolate components that may or may not be present in a given chimera. As an example, the above mentioned chimeric regulator SLCP_GL_ ^25^ is identified as LacI-GGBP-OD.

### Novel multiplexed assembly to create the library of transcriptional regulators

The construction of thousands of chimeric TFs by gene fusion required designing a systematic pipeline. A general outline of the assembly strategy employed for the construction of the libraries of TFs can be found on *Figure 2*. Briefly, we planned a set of 4275 “core chimeras” (15 DBDs x 19 LNKs x 15 SBPs). Beyond the “core chimeras” the total number of TFs that could potentially be generated is the result of assembling 119 DBD-ENDS, 19 LNK/no LNK, 30 Ct-SBD and 2 OD/no OD combinations, totalling 135,660 chimeras. To simplify the synthesis of all of these fusion genes they were assembled without OD and subsequently cloned in two expression vectors, pCKTRBS and pCKTRBS-OD (*Methods*, *Table S2*). The TFs cloned into pCKTRBS-OD incorporated a 3’ translational fusion to a OD sequence included in the plasmid chassis, whereas the TFs cloned into pCKTRBS did not. Cloning the genes in this fashion generated two libraries, each containing 67,830 chimeras.

**Figure 2.**
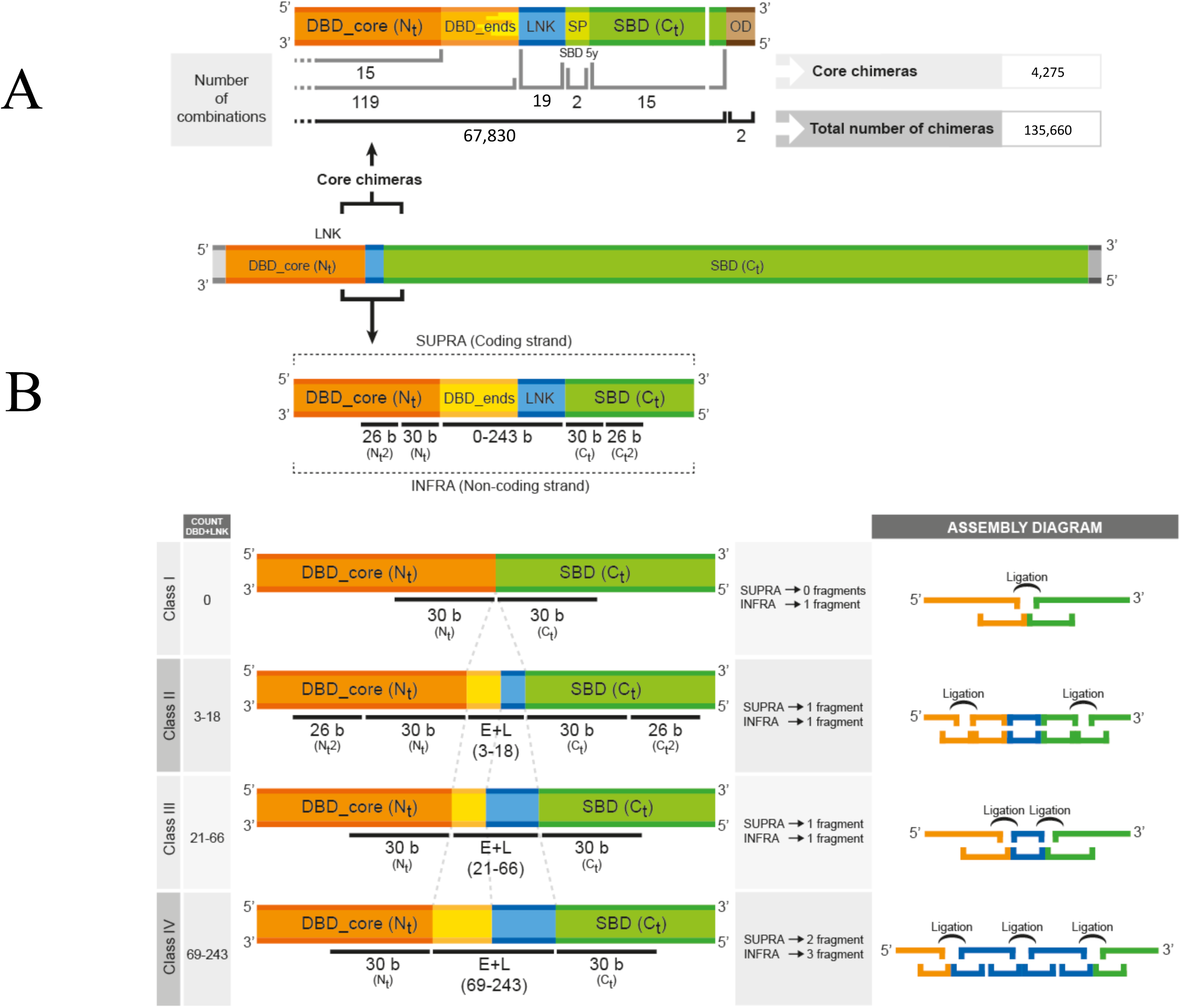
Representation of the domain structure of the chimeric TFs and their classification depending on the complexity of their assembly via eLCR. Representation of an average chimera maintaining the proportion between the lengths of the different regions that integrate it. DBD-CORE are represented in orange, ENDS are represented in dark yellow, LNK in blue, signal peptides (SP) in light green, SBD in dark green and OD in brown. **A)** Not-to-scale representation of the joint between DBDs and SBDs together with OD. The number of domains of each kind incorporated into the assembly scheme as well as the maximum number of chimeras resulting from the eLCR are also represented. **B)** Zoomed in simplified version of the joint between DBDs and SBDs. ENDS and LNKs occupy the center of the representation, as they were the regions incorporated in the eLCR through a collection of oligonucleotides. DBDs and SBDs (with and without SP) were introduced in the assembly as dsDNA fragments. The numbers below the different domains represent the number of DNA bases included into the oligonucleotides in the DNA microarray synthesis. The panel below represents the number of bases corresponding to each one of the domain types that needed to be introduced into the eLCR oligonucleotides depending on the assembly Class the chimera was assigned to. The assembly diagram to the right represents, in the same color code that above, the final *Supra* and *Infra* oligonucleotides and the ligation events that needed to take place to reconstitute the final chimera.

To connect Nt-DBD and Ct-SBD domains with a diverse set of LNK sequences, we developed a new cloning strategy based on the LCR (Ligase Cycling/Chain Reaction) assembly method ^46^. We refer to this new method as “enhanced LCR” or eLCR for short (*Methods*). Conventional LCR provides scarless fusion of dsDNA fragments. However, eLCR allows the targeted insertion of oligonucleotides in the junction between those fragments, enabling the generation of many different connections between DBD and SBD domains. Every DBD in the library was composed of a fixed sequence identified as DBD-CORE that had to be connected to the SBD. These two domains were bridged by two elements: i) specific extensions of every DBD-CORE called ENDS; ii) LNK sequences. eLCR enabled the use of oligonucleotides to attach all the possible ENDS to their correspondent DBD-CORE plus a LNK option per chimera. *Figure 2* is a schematic representation of the different classes of chimeras present in the library attending to the number of oligonucleotides necessary to assemble them. The rationale behind this design is explained in detail in *Methods* and *Supplementary Materials*. The fusion genes assembled by eLCR were PCR amplified with oligonucleotides that incorporated adapter sequences homologous to the expression vectors pCKTRBS or pCKTRBS-OD (*Table S2*). The adapter to pCKTRBS-OD created a translational fusion to the LacI OD integrated in the vector. All the oligonucleotides necessary for these amplifications are listed in *Table S3*. The libraries of bacteria expressing the collections of chimeras from pCKTRBS/pCKTRBS-OD were named *E. coli* (Ch-END) and *E. coli* (Ch-OD) (*Table S2*)

### Design and construction of synthetic reporters for screening the library of transcriptional regulators

The reporter system designed to screen the activity of the libraries of chimeras was based on the controlled expression of GFP by synthetic promoters. In order to assay the activity of a given chimera, a reporter plasmid specific for the DBD of said chimera was needed. Having 15 DBD modules in the assembly mix, we constructed 14 modified *Ptac* promoters (the extra one to 15 was *Ptac* itself), each one engineered to contain operator boxes for a particular DBD. The sequence of these 15 promoters is included in (*Table S4*). The high-copy number family of reporter vectors pHC_DYO*DBD*-R (*Table S2*) was constructed by integrating the above mentioned 15 promoters into a pUC chassis making them drive the expression of GFP (*Methods*). In the presence of chimeric TFs that retain their DNA-binding ability, the expression of the GFP reporter gene should be reduced. On the other hand, when the inducer benzoate molecule is added to the culture, the repression of the reporter should be relieved only where the chimeras are functional.

The screening design based on the enrichment of benzoate-binding chimeras from a complex pool required the synthetic promoters integrated into pHC_DYO*DBD*-R plasmids to show a basal expression as similar as possible. The objective was to minimize the intrinsic biases of the pipeline towards the enrichment in TFs whose DBD interacted with the strongest promoters. Several versions of the modified promoters were created and inserted into pHC_DYO*DBD*-R. Their basal expression levels compared to *Ptac* were empirically assessed *in vivo* (*Methods*). The ones whose activity fell outside a threshold of one order of magnitude were discarded. *Figure S1* shows the relative level of basal expression of the final set of synthetic promoters compared to *Ptac*. As a complement to this individual assessment of activity a mix of strains carrying all the reporter promoters was assayed in FACS, demonstrating that once the strain carrying the wild-type promoter *Ptac* was removed from the mix the rest of strains were not preferentially enriched based on their basal GFP expression (*Methods*). The collection of reporter plasmids was transformed on *E. coli* (Ch-END) and *E. coli* (Ch-OD) as described in *Methods* The resulting libraries in which every bacterium was expressing a chimera and carrying a reporter promoter were identified as AYC Lib-Ch-END and AYC Lib-Ch-OD (*Table S2*).

### Functional screening of the library of transcriptional regulators using the synthetic reporters

Screening was based on two-step enrichment as represented in *Figure 3*. In the first step (Negative Sorting), the population was sorted to enrich the cells in which there was a pairing between the DBD of the chimera and the operator boxes driving GFP expression in the reporter promoters. In these cells, the chimeric TF binds to the reporter promoter repressing the expression of GFP. As the negative selection was a functional screening its benefits went beyond recovering a sorted population enriched in properly paired chimera-reporter plasmids. Bacteria with the correct pairing of TF and reporter but where the chimeric regulator was not a functional repressor (e.g. due to wrong protein folding or lack of secondary modifications in the polypeptide) were discarded. In a second step (Positive Sorting) the bacteria recovered after the Negative Sorting were cultured in the presence of the inducer molecule benzoate then sorted for expression of GFP. Bacteria expressing functional chimeras required two events to be recovered: a) their SBD recognized benzoate; b) the conformational change experimented when this happened could alter the allostery of the repressor forcing it to unbind from its DNA operator in the reporter promoter. In the cells where these two events happened there was no obstacle for RNA polymerase-σ^70^ to recognize the -10/-35 of the promoter and transcribe GFP, increasing the fluorescence of the bacteria. FACS sorting was programmed to recover the most fluorescent cells in the GFP emission range. The positive sorting was repeated to increase the enrichment.

**Figure 3.**
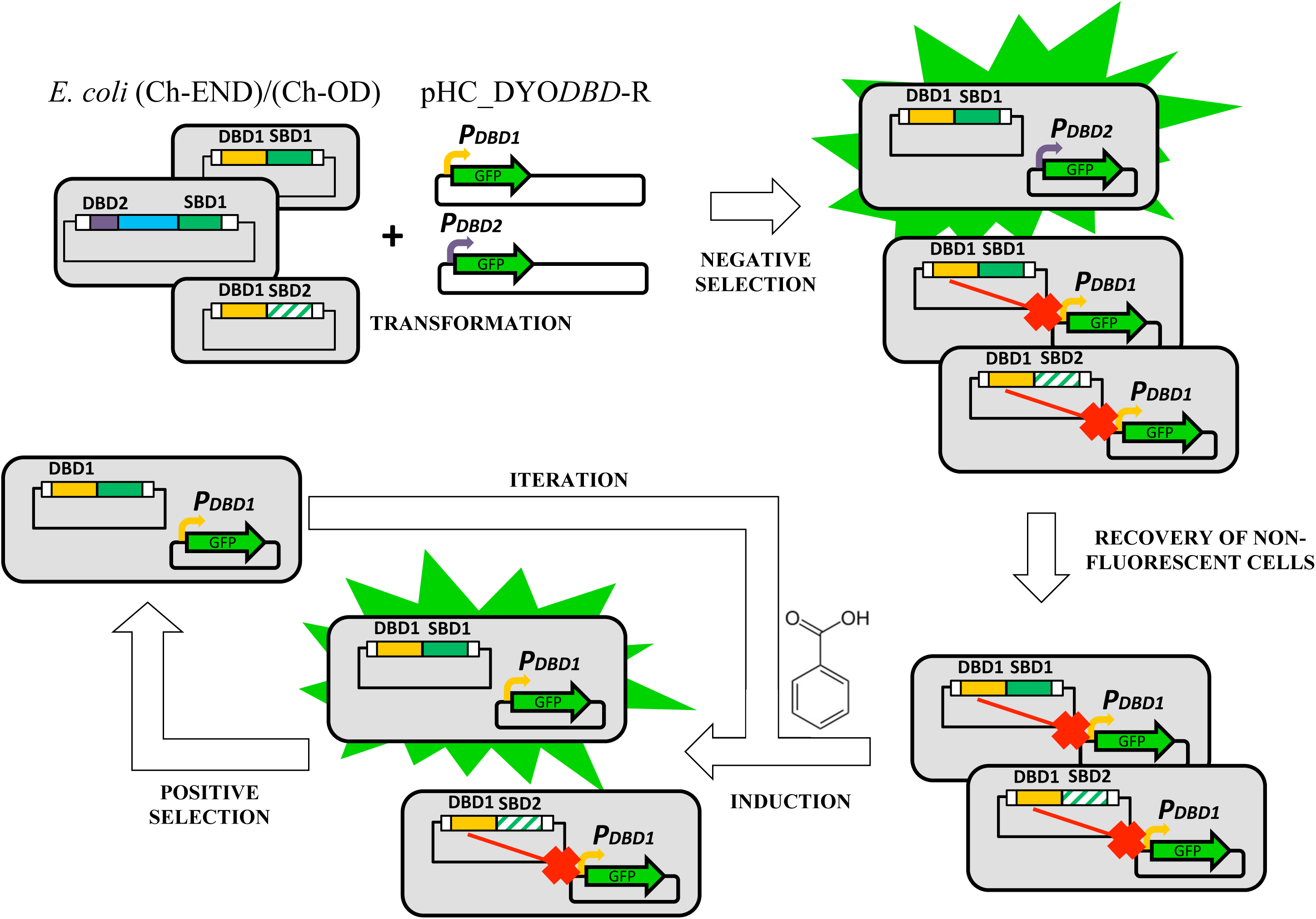
Schematic representation of the enrichment process. Chemically competent *E. coli* (Ch-END)/(Ch-OD) cells (grey rectangles) carrying vectors pCKTRBS/pCKTRBS-OD expressing chimeric TFs were transformed with reporter plasmids pHC_DYO*DBD*-R as described in *Methods*. The resulting cells were grown in LB media in the presence of aTc. Under this culture conditions the chimeric TFs were expressed. Every TF contains one out of 15 DBD. In the cells where the DBD of the chimera (represented as yellow and purple rectangles) was able to recognize the operator boxes of the reporter promoter (bent arrows in yellow and purple) introduced in pHC_DYO*DBD*-R, and it retained its DNA-binding capabilities, the expression of GFP is repressed. On the other hand, GFP was highly transcribed in the cells where the TF was not able to interact with the reporter promoter. The cells showing the lowest levels of GFP expression were recovered using the FACS enrichment described in *Methods* in the so called Negative Sorting. The recovered cells were grown in the presence of the inducer molecule (benzoate). TFs carrying a functional SBD were able to recognize the inducer and transduce that information to the DBD triggering a conformational change strong enough to detach the DBD from the DNA. Under these conditions bacteria carrying functional TFs would resume the transcription of GFP from the reporter promoter. A FACS sorting allowed to obtain a population enriched in the functional chimeras (Positive Sorting). This enrichment process was iterated.

The viability of using FACS for our enrichments was confirmed by performing control sortings as described in *Supplementary Materials*. Initially, we recovered a functional wild-type regulator, LacI, from a library of TFs. In a second phase we used the glucose-inducible chimera LacI-GGBP-OD (SLCP_GL_) as our reference strain, since it is one of the few published chimeric transcriptional repressors ^47^. LacI-GGBP-OD was dramatically more abundant in our libraries after enriching it for glucose-induced TFs, both when this chimera was added to the pool exogenously and when it was assembled as another fusion gene in the library together with the benzoate-binding chimeras. For the screening for benzoate-responsive chimeras, AYC Lib-Ch-END and AYC Lib-Ch-OD were grown separately in LB supplemented with aTc. Non-fluorescent cells plus the 10% of fluorescent cells with the lowest fluorescence were sorted. The recovered cells were grown in the presence of aTc and benzoate (chimeras expressed, benzoate-responsive TFs allowing GFP expression) and subjected to several enrichment cycles in which the most fluorescent cells were recovered every time. *Table S5* summarizes the percentages of cells recovered for both AYC Lib-Ch-END and AYC Lib-Ch-OD libraries in every step of the enriching process. AYC Lib-Ch-OD was subjected to an extra cycle of enrichment, since the improvement of fluorescent construction after every iteration seemed to be more increased than that of AYC Lib-Ch-END. The distribution of TFs in the starting libraries and the ones recovered after 4-5 cycles of enrichment was analyzed by next generation sequencing (*Methods*). There was not a single chimera dominating the population,but on average 33 chimeras were present in the enriched libraries in an abundance equal or superior to 1%. These results suggest that in the quest to find a benzoate-responsive chimera in the tested libraries there was not a single optimal solution but an array of chimeric TFs that were functional to some extent. This was the expected outcome of the library analysis, since the chimeras assembled in this work have not been subjected to the continuous forces of evolution natural TFs have experienced.

### Validation of select benzoate-sensing chimeric regulators

Of the more than hundred chimeras that showed enrichment in the screening, two were selected for further characterization because of their unique features. First, from the AYC Lic-Ch-OD library we chose CbnR-ABE44898-OD (ChTFBz01) because it carried a Ct LacI OD and its SBD (ABE44898) was similar to RPA0668, a *R. palustris* protein whose ability to bind benzoate had already been tested ^48–50^. It is important to note that even though ABE44898 was classified as a PBP, its SP (included in this construction) sequence was the most divergent among all SBDs. This peculiarity may have had conferred interesting biophysical properties to the chimera, since the SP could be working as an extended linker sequence. Second, from the AYC Lib-Ch-END library we chose LmrR-BzdB1_nSP (ChTFBz02), containing as SBD a PBP associated to a cluster responsible for benzoate catabolism ^51,52^. Preliminary analysis suggested BzdB1 is directly involved in the binding of benzoate and is highly soluble (unpublished data, J.F. Juarez and E. Diaz). The inclusion of proteins like BzdB1, associated to clusters involved in the metabolism of small soluble molecules allowed for the improvement of the libraries by the addition of domains presenting higher chances to bind the inducer molecule. The lack of SP in the chimera was likely to reduce the flexibility of the DBD-SBD interface, which might be positive for certain DBD-SBD combinations.

ChTFBz01 and ChTFBz02 were respectively re-cloned in pCKTRBS-OD and pCKTRBS. Afterwards, the strains carrying them were transformed with the appropriate reporter vectors. The resulting strains AYC ChTFBz01 and AYC ChTFBz02 (*Table S2*) allowed us to assay *in vivo* the ability of CbnR-ABE44898-OD and LmrR-BzdB1_nSP to repress the expression of GFP and be de-repressed by benzoate. Following the experimental design described in *Methods* we found that, in both cases, there was a reduction of the GFP fluorescence of the culture when the chimeras were expressed in regular culture medium (chimeras expressed, GFP promoter repressed). However, when the inducer benzoate was added to the cultures the strains showed GFP levels very close to the basal expression of the reporter promoters. The fold difference of GFP expression between the repressed and induced conditions is 3.06 ± 0.74 for CbnR-ABE44898-OD and 3.13 ± 0.31 for LmrR-BzdB1_nSP. *Figure 4* shows the expression levels of the reporter promoters of the assayed constructions when the chimeric repressors were expressed. As we can observe the addition of benzoate restored expression levels close to the basal ones, suggesting that both CbnR-ABE44898-OD and LmrR-BzdB1_nSP were functional sensors able to repress the expression of their target promoters and cease said repression in the presence of their inducer molecule, benzoate. A detailed *in vitro* study of the interaction between the purified chimeras and the reporters would be needed to calculate the constants driving the kinetic parameters of repressor-promoter and repressor-inducer interactions, but the evidence present in this work shows these synthetic regulatory circuits worked *in vivo*. It is important to note that there is vast room for improvement of these new synthetic regulators using technologies such as directed evolution and semi-rational protein design ^53^. The point where these TFs are now would be equivalent to the point in evolution history where LacI was when its DBD recruited the ancestor of its SBD domain ^24^, millions of years prior to genetic evolution optimizing the communication between the domains that shaped that ancestral chimera into the TF we know today.

**Figure 4.**
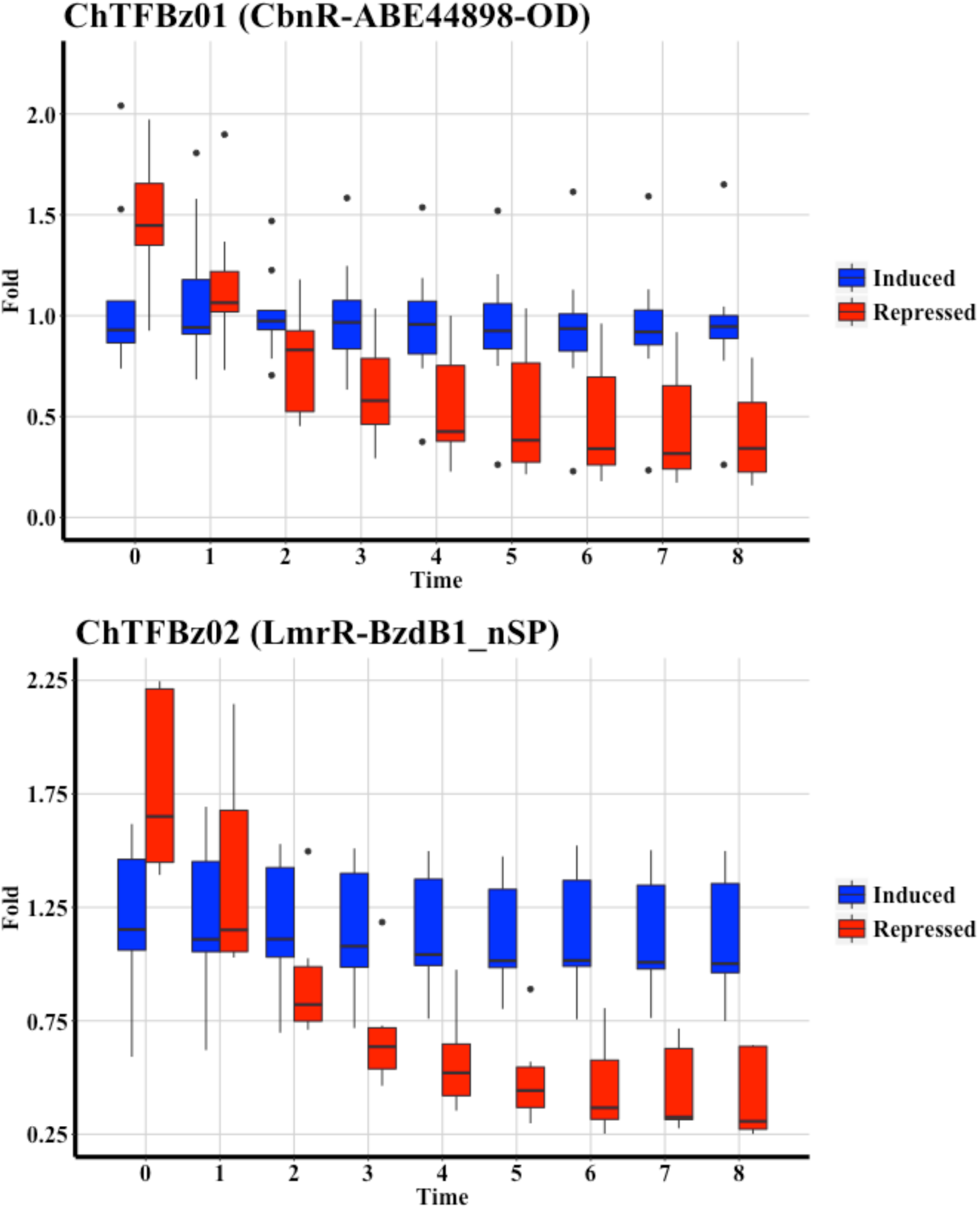
*In vivo* demonstration of the functionality of the ChTFBz01 (CbnR-ABE44898-OD) and ChTFBz02 (LmrR-BzdB1_nSP) chimeras. Time course showing relative GFP fluorescence on AYC ChTFBz01 and AYC ChTFBz02 strains grown in minimal medium in a multi-well plate reader as indicated in *Methods*. Promoter activity is measured as fold of the relative fluorescence (Fluorescence in arbitrary units / OD600) of the strains grown under inducing conditions (cultures supplemented with aTc and benzoate; blue boxes) or repressing conditions (cultures supplemented with aTc; red boxes) compared to the basal expression of the promoter (no aTc, no benzoate). It can be appreciated how in order to maintain the basal activity (Fold=1) the addition of benzoate to the culture medium is necessary when the chimeras are expressed by the addition of aTc. The difference between repressed and induced expression is statistically significant from the third hour of the culture on. Growth conditions and fluorescence assays performed as described in *Methods* (n=6-10).

The chimeragenesis method described in this study is based on the hypothesis that most TFs resulting from the assembly would be either nonfunctional or toxic for the host cells ^54^. One of the main advantages of our FACS enrichment system is that it assays the functionality for every construction in the competitive environment of a bacterial library. Toxic and poorly functional constructions can be weeded out in successive enrichment steps that are tuned to be more stringent in every step. The utility of the pipeline presented in this work has been demonstrated by obtaining biosensors for benzoate. Even though we have focused our efforts on obtaining benzoate-responsive TFs, our library was designed as well to generate a transcriptional repressor induced by 4-hydroxybenzoate. Both benzoate and 4-hydroxybenzoate are environmentally relevant, due to their association to the degradation of lignin ^50,55,56^. Their abundance makes them a principal target in the valorization of lignin and its associated aromatic monomers ^57^. To the best of our knowledge there are no published transcriptional repressors directly induced by benzoate. The catabolism of this aromatic molecule has been thoroughly studied in anaerobic pathways, where the most frequent inducer for repressors associated to these clusters is the first metabolite of the pathway benzoyl-CoA ^58^, requiring the attaching of subsidiary benzoyl-CoA ligase enzymes ^51,59^ to the repressor-promoter system in order to apply them to synthetic regulatory circuits. BzdR ^59^ and BoxR ^59^, two well-studied regulators associated with benzoate catabolism in anoxic conditions indeed respond to benzoyl-CoA and not to benzoate. It is unknown whether benzoate or benzoyl-CoA are the actual inducers of one and two-component systems repressing catalytic pathways related with benzoate in different microorganisms, e.g. BamVW in *Geobacter metallireducens* ^60^, BadM in *Rhodopseudomonas palustris* ^61^ and BgeR in *Geobacter bemidjiensis* ^62^. BenM, a transcriptional activator able to recognize benzoate as an inducer molecule has been extensively studied ^34,63,64^. Despite the existence of this activator we still consider that a benzoate-responsive transcriptional repressor has added advantages. The TFs we have constructed in our libraries base their function in the the simplest transcriptional repression mechanism, interference with RNA-polymerase sigma factor binding to DNA ^65^. Moreover, the chimeric genes presented in this work present a single binding pocket for a small molecule, this is, the binding site of the PBP domain, whereas BenM can simultaneously bind two inducers (benzoate and *cis*,*cis*-muconate) ^33^ creating more complicated inducer landscapes and potentially undesired unspecific activations not dependent on benzoate.

### Conclusions

In this study, we describe a comprehensive strategy to create custom monogenic biosensors using fusion of modular components. Using this strategy, we constructed multiple novel benzoate-sensing TFs and further characterized two: ChTFBz01 and ChTFBz02. These novel sensor proteins expand the limited collection of available transcriptional repressors that can be used as biosensors for the degradation of lignin. Beyond their immediate usability to tackle this biotechnological problem, they represent an important milestone for the construction of any TF on demand since our pipeline can be readily applied to create many other custom-made chimeric TFs. Using periplasmic binding proteins as detection domains underscores the potential of this method in generating tailored biosensors far beyond swapping of domains between already known regulators.

The construction of a large number of fusion genes by assembling different collections of modules in an appropriate order was made possible by the development of eLCR, a novel strategy to assemble highly diverse fusion libraries. presented in this work. While preserving the ability to attach DNA fragments in a scarless fashion this assembly enables incorporating short DNA sequences between said fragments. This characteristic makes this procedure ideal for constructing libraries of mutant proteins that carry, for example, linkers of different lengths or clusters of mutations. Another strength of this method resides in its potential to create novel TFs through a straightforward design that does not demand advanced knowledge of protein structure to succeed.

## METHODS

### Bacterial Strains, Plasmids, and Growth Conditions

Bacterial strains and plasmids used are listed in *Table S2*. *Escherichia coli* strains were grown in lysogeny broth (LB) medium ^66,67^ at 37°C. When required, *E. coli* cells were grown in M63 minimal medium ^68^ using the corresponding necessary nutritional supplements and 30 mM glycerol (Sigma, St. Louis, MO), as carbon source. When appropriate, antibiotics were added at the following concentrations: ampicillin/carbenicillin (100 μg/ml), chloramphenicol (25 μg/ml) (Sigma, St. Louis, MO). When the expression of protein from pCKTRBS/pCKTRBS-OD was required the cultures were induced with 0.5 μg/ml anhydrotetracycline (aTc) (Clontech, Mountain View, CA). 1 mM IPTG (Sigma, St. Louis, MO), 1mM sodium benzoate (Sigma, St. Louis, MO) and 0.4% glucose (Teknova, Hollister, CA) were added to the media when the experimental conditions required it.

### Molecular Biology Techniques

Standard molecular biology techniques were performed as described previously ^69^. Plasmid DNA was purified with a Quiaprep Spin Miniprep Kit (Qiagen, Hilden, Germany). DNA fragments were purified with DNA Clean-up & Concentration Kit (Zymo research, Irvine, CA). The oligonucleotides employed for PCR amplification of the cloned fragments and other molecular biology techniques are summarized in *Table S3* and were supplied by IDT (Coralville, IA). All cloned inserts and DNA fragments were confirmed by Sanger sequencing ^70^ performed by Genewiz Inc. (Cambridge, MA). Commercially available *E. coli* NEB5-alpha and *E. coli* NEB5-alpha *F’ I*^*q*^ chemically competent cells (NEB, Ipswich, MA) were used for routine transformations. Alternatively electrocompetent *E. coli* cells were generated and used immediately (Gene Pulser; Bio-Rad, Hercules, CA) ^69^. Cloning was routinely performed by Gibson assembly ^71^. Nucleotide sequence analyses were done at the National Center for Biotechnology Information (NCBI) server (http://www.ncbi.nlm.nih.gov). Prediction of signal peptides was performed using SignalP ^72^ at the Technical University of Denmark online server (http://www.cbs.dtu.dk/services/SignalP/).

### Construction of the expression vectors pCKTRBS and pCKTRBS-OD

The plasmid pCK01 ^73^ was the chassis of choice for the construction of pCKTRBS. This vector was tasked with the expression of all the chimeric TFs generated in this work. The *Plac* promoter of pCK01 was replaced by a *tetR-PtetO* cassette modified from the plasmid pKD154 (B.L. Wanner and K. Datsenko, unpublished). *PtetO* would be the inducible promoter transcribing the chimeras. In the absence of aTc *PtetO* was repressed by TetR and the chimera did not get expressed, whereas in the presence of aTc TetR unbound *PtetO*, opening the system and enabling transcription from it. The resulting pCK01 derivative was furtherly modified to include a consensus RBS sequence downstream of *PtetO*, hence the name pCKTRBS (“T” for t*etR*, RBS for the consensus Shine-Dalgarno). The RBS and the adjacent upstream bases constituted a region to which the 5’ homology arm included in all the DBD domains held homology. A fragment of the old pCK01 polylinker located at 3’ of the newly inserted RBS was used as the homology arm at 3’ of the SBD that were amplified to be cloned without further modification. A variant of pCKTRBS containing LacI OD was constructed introducing OD between the consensus RBS and the remains of the former pCK01 polylinker. The amplification of the chimeras with a 3’ arm homolog to LacI OD granted the cloning of the TFs as a translational fusion to OD in their Ct end. This new vector was named pCKTRBS-OD. Since pCKTRBS contained a constitutive promoter driving the expression of TetR, the expression of the genes cloned in pCKTRBS/pCKTRBS-OD was repressed by default. In the cases where it was necessary to induce the expression of the cloned gene aTc was added to the culture media as indicated above. A detailed representation of the region comprising from *tetR* to OD in pCKTRBS-OD including operator boxes for TetR and the RNA polymerase-σ^70^ subunit located on the expression promoter *PtetO* is shown in *Figure S2*. The fully annotated sequence of pCKTRBS and pCKTRBS-OD is shown in *Supplementary Materials*.

### Synthesis, amplification and purification of dsDNA and ssDNA fragments for eLCR

DBD-CORE and SBD domains were synthesized as dsDNA fragments by the custom DNA manufacturer Gen9 (Cambridge, MA). Silent mutations were incorporated as requested by the manufacturer when it was necessary to remove target sites for restriction enzymes that would be needed for the assembly process.

The sequence of all the possible DBD-CORE-ENDS-(LNK)-SBD combinations as well as a list containing all the oligonucleotides necessary to assemble them was obtained using a custom Perl script. Two types of oligonucleotides were employed in the assembly depending on their orientation and the DNA strand they would conform when ligated. The ones identified as *Infra* had the same sequence and orientation that the template strand, whereas the ones labeled as *Supra* carried the same sequence and orientation that the coding strand (*Figure 2*). Specific adapters for *Infra* and *Supra* were engineered based on two orthogonal pairs of primers who had been designed by ^74^ to selectively amplify pools of oligonucleotides from high-fidelity DNA microchips. These adapters were 5’ and 3’ flanking sequences already incorporated into the oligonucleotide synthesis order (*Figure S3-A*) Three Custom Array chips were used to synthesize the 223170 required oligonucleotides. DNA synthesis in microarrays was performed by Daniel Wiegand in the Synthetic Biology Platform from the Wyss Institute at Harvard (Boston, MA). Oligonucleotide libraries were synthesized using standard phosphoramidite chemistry on a 90K CustomArray DNA Oligonucleotide Library Synthesis (OLS) microarray using the B3 Synthesizer platform. After libraries were synthesized, the surface bound oligonucleotides were cleaved from the microarray by incubating them in 30% ammonium hydroxide at 65°C for 12 hours. Oligonucleotide libraries were then dried and concentrated with a SpeedVac Concentrator set at 65°C and vacuum engaged for 3 hours. The resulting pellet was resuspended in 70 uL of 1X TE buffer and purified using a P-30 size exclusion column (Biorad) pre-equilibrated with 1X TE buffer. The concentration of the resulting purified oligonucleotide libraries was determined with a NanoDrop spectrophotometer at A260. The ssDNA pool resulting from the synthesis was amplified to preserve the original sample and to enable the obtention of a more concentrated suspension of oligonucleotides.

*Figure S3* is an schematic representation of the adapters enabling the amplification of the oligonucleotide library (Panel A) as well as of the purification process (Panel B). The pool of oligonucleotides obtained from the microarray synthesis was PCR amplified in two different reactions. The primer pair Lib_Adapt_Supra_5 / Lib_Adapt_Supra_3 amplified specifically *Supra* ssDNA, whereas Lib_Adapt_Infra_5 / Lib_Adapt_Infra_3 amplified *Infra* oligonucleotides. Lib_Adapt_Supra_5 and Lib_Adapt_Infra_5 oligonucleotides used for the PCR reaction carried two modifications: i) a biotin in the 5’-end; ii) a uracil instead of a timidin in their 3’-end. These modified versions of the oligonucleotides were identified as BioT-Lib_Adapt_Supra_5-U and BioT-Lib_Adapt_Infra_5-U. PCR amplification of the oligonucleotide libraries using the modification-bearing oligonucleotides as primers generated biotinilated and uracilated dsDNA fragments. Once the CustomArray DNA pools had been amplified it was necessary to remove the adapter sequences that enabled the hybridization of Lib_Adapt_Supra_5 /Lib_Adapt_Supra_3 and Lib_Adapt_Infra_5 / Lib_Adapt_Infra_3. Type IIS enzymes were used in a first step for the removal of the non-biotinilated end thanks to their ability to cut dsDNA bases away from their recognition boxes in the target DNA. The adapters where Lib_Adapt_Supra_3 and Lib_Adapt_Infra_3 hybridized incorporated BsmFI and BspQI target sequences, respectively. dsDNA *Supra* fragments were digested with BsmFI and the *Infra* with BspQI, following in both cases the specifications of the manufacturer (NEB, Ipswich, MA). Digested pools were incubated with Dynabeads M-280 Streptavidin (Life Technologies, Carslbad, CA). The dsDNA oligonucleotides got attached to the paramagnetic beads through the biotin and the adapters cut in the above mentioned digestion were washed away. Non-biotinylated strands were not directly attached to the beads and could be removed by an alkaline denaturalization in 125 mM NaOH at 90°C for 2 minutes and, afterwards, washed away. The *Supra* and *Infra* oligonucleotides were detached from the beads and removed from the remaining adapter through USER digestion (NEB, Ipswich, MA), that cut the DNA in the uracil incorporated at the 3’-end of the biotinylated primers. The final ssDNA *Supra* and *Infra* pools were purified using DNA Clean-up & Concentration Kit (Zymo, Irvine, CA).

### eLCR (Enhanced LCR)

Canonical LCR is based in the following principle: short ssDNA fragments with complementarity to two DNA molecules that are going to be assembled act as staples to locate them in close vicinity allowing for a DNA ligase to bind them ^75,76^. A initial denaturation at high temperature separates the two strands of the target DNA fragments. When the temperature decreases the staple oligonucleotide anneals to the target sequences and the nick between the two DNA molecules is closed by a thermostable DNA ligase in a scarless process. This assembled strand serves as template for the complementary strand in a subsequent cycle of denaturalization, annealing and ligation. Temperature cycling allows the assembly of multiple DNA fragments in a highly efficient manner. We confronted a major challenge when trying to apply the original LCR method to the assembly of the fusion genes presented in this work. Different sequences needed to be introduced between DBD and SBD, bridging them, on top of the scarless assembly provided by LCR. Between the 15 DBD-CORE and the 30 SBD (counting versions with and without their SP) it was needed to add the specific ENDS corresponding to each DBD-CORE and the 19 different LNK options. The length of the sequences introduced between the DBD-CORE and the SBD ranged between 3 bp (shorter DBD-END equivalent to one codon) and 244 bp (combining the longest DBD-END, 120 bp, and the longest LNK, 123 bp). We created four different classes of chimeras based on the the number and complexity of the stapling oligonucleotides needed to assemble them, which was related to the length of the sequence introduced in between DBD-CORE and SBD. Class I chimeras (1 *Infra*) were TFs in which there was a direct connection between DBD and SBD. More than half of the library belonged to Classes II (1 long *Infra*,1 *Supra*) and III (1 short *Infra*,1 *Supra*), chimeras in which 3 to 66 bp were introduced between DBD and SBD. Finally, Class IV (2 *Infra*,3 *Supra*) chimeras carried a bigger than 66 bp insertion. More information about the different classes of chimeric TFs can be found on *Supplementary Materials*.

The first step for the eLCR assembly consisted on the phosphorylation of all the DNA molecules taking part on it ^77^. An equimolar mix of the 15 DBD-CORE and 30 SBD was mixed with the product of the amplification of *Supra* and *Infra* oligonucleotides and phosphorylated using T4 Polynucleotide Kinase (Enzymatics, Beverly, MA) for 30 minutes at 37°C. These phosphorylated products were the substrate for the assembly of the chimeric TFs using a thermostable DNA ligase. Prior to the assembly of the complete collection of TFs we tested a mock library of four evenly distributed chimeras, each one of them belonging to one of the classes described above. DBD and SBD of this testing library were taken from the collection of DNA fragments used for the final construction of the chimeras, whereas the subset of oligonucleotides necessary to assemble them was provided by IDT (Coralville, IA). This mock library enabled the testing of three different thermostable ligases: Ampligase (Epicentre, Madison, WI), 9°N DNA ligase (NEB, Ipswich, MA) and *Taq* DNA ligase (NEB, Ipswich, MA). After this preliminary optimization *Taq* DNA ligase seemed to be the most efficient ligase for our new eLCR technique. Another parameter that needed to be optimized with the mock library was the thermal treatment of the reaction mix once the thermostable DNA was added to it. The only chimeras that were generated through a canonical LCR were Class I, since they were assembled exactly as the chimeras described in ^77^. Temperature cycling allows a ligated strand to serve as a template for the single ligation required to generate its reverse-complementary strand, as is the case for Class I chimeras. However, the assembly of Classes II, III and IV required more events to happen, since several oligonucleotides needed to be hybridized and ligated to reconstitute the DNA strands. We found that the cycling of temperatures helped to increase the proportion of Classes II, III and IV in the pool. The relative distribution of the different classes of chimeras in the mock library was affected by the cycling program of choice. The final thermal treatment for the assembly was the one used by ^78^: an initial incubation at 94°C for 2 minutes is followed by 10 cycles of alternating 94°C for 30 seconds and 45°C for 4 minutes. In order to facilitate the proper hybridization of the staple oligonucleotides the speed of the temperature ramp decrease from 94°C to 45°C was reduced to a 30% of the top speed in the Mastercycler Pro (Eppendorf, Hamburg, Germany) thermocycler used for the assay. The products of the assembly were PCR amplified with primers hybridizing on the flanking sequences introduced to 5’ of the DBD and 3’ of SBD. The primer RBS_Tail_F1 hybridized in the 5’ 30-b region common to all the DBD domains. The primers END_Tail_R1 and OD_Tail_R1 hybridized in the 30-b regions designed to clone the chimeras without or with a translational fusion to OD, respectively (*Table S3*). The amplification was monitored in real time using a LightCycler 96 System (Roche, Basel, Switzerland) and stopped in mid-exponential phase of the amplification curve, minimizing non-quantitative effects in the plateau of the reaction that would contribute to reduce the diversity of the library ^79^.

### Cloning of eLCR-assembled genes

Preliminary data suggested that an intermediate cloning of the amplified library using a Zero Blunt TOPO PCR cloning Kit vector (Invitrogen, Carlsbad, CA) instead of a direct cloning into pCKTRBS/pCKTRBS-OD was more efficient. The purified products of the PCR amplification of the libraries, with and without OD, were cloned into pCR-BluntII-TOPO following manufacturer's instructions and transformed into chemically competent NEB5-alpha *F’ I*^*q*^ cells (NEB, Ipswich, MA). An aliquot of the recovered cells was stored at -80°C to preserve the maximum possible diversity of the TF collection. Another aliquot was used to isolate plasmids. These plasmid libraries were the template for new PCR amplifications of the chimeras (monitored in real time as described above). One amplification batch included all the chimeras carrying OD (using the primer pair RBS_Tail_F1 / OD_Tail_R1) and another batch included the TFs without OD (using the primer pair RBS_Tail_F1 / END_Tail_F1). The amplified libraries were cloned into pCKTRBS/pCKTRBS-OD vectors using Gibson assembly. Linear pCKTRBS/pCKTRBS-OD vectors were obtained by divergent PCR amplification using the primer pairs pCKTRBS_RBS_Tail_R1 / pCKTRBS_END_Tail_F1 and pCKTRBS_RBS_Tail_R1 / pCKTRBS-OD_OD_Tail F1, respectively (*Table S3*). The 30-bp 5’ and 3’ terminal regions of both plasmids showed homology to the sequences flanking the assembled chimeras, so that when they got integrated they were located downstream the *PtetO* promoter and the RBS. The products of the library assembly into the expression vectors pCKTRBS/pCKTRBS-OD were transformed into chemically competent NEB5-alpha cells (NEB, Ipswich, MA). The resulting collections of bacteria carrying chimeric TF cloned into pCKTRBS/pCKTRBS-OD were named *E. coli* (Ch-END) and *E. coli* (Ch-OD) respectively.

### Construction of the pHC_DYO*DBD*-R library of reporter vectors

A set of reporter vectors designed to monitor the activity of the libraries of TFs was constructed on a pUC19 chassis ^80^. Preliminary data revealed that the expression of GFP from high-copy plasmids versus low-copy plasmids was more suited to monitor the activity of the regulators using FACS. Plasmid pHC_DYOLacI-R (*Table S2*) was the first one constructed and tested. The full sequence of pHC_DYOLacI-R and the origin of the different regions that were PCR amplified and Gibson-assembled to build it are detailed in *Supplementary Materials*. pHC_DYOLacI-R carried a *Ptac* promoter driving the expression of the reporter gene GFP. The *Ptac* included in this first reporter plasmid was the scaffold on top of whom the rest of the synthetic promoters were built (*Table S4*). One of the main requisites to select the transcriptional repressors donating their DBDs to the chimeric TFs was that they had their operator boxes published and experimentally tested (*Table S1A*) so that they could be placed in synthetic promoters driving the expression of the reporter gene and thus being used to screen TFs binding to them. *Ptac* was modified to include the operator boxes for the rest of the DBDs preserving the integrity of the -10/-35 boxes so that they could still be recognized by the RNA polymerase-σ^70^ subunit. The distance between -10/-35 was maintained constant for the same reason. The operator boxes published for every regulator were placed on *Ptac*. They were strategically located so that when the protein that recognized them was bound to the DNA transcription from the modified promoter would cease. For this purpose location within the spaces between -10/-35 and between the transcription start site and -10 were preferred. In some occasions mutations in -10/-35 were required to accommodate the operators. *Table S4* contains the sequences of every synthetic promoter. The method of choice to construct the pHC_DYO*DBD*-R vectors consisted on divergent PCR using pHC_DYOLacI-R as template (see oligonucleotides in *Table S3*). The divergent oligonucleotides amplified the whole extent of pHC_DYOLacI-R but for the *Ptac* promoter and LacI operator boxes. There was a devoted pair of primers for every reporter plasmid that was constructed. The modified *Ptac* sequence for every given synthetic promoter was split between the 5’ (phosphorylated) and 3’ oligonucleotides. The PCR products were amplified and ligated so that they would close within themselves, restoring in the joint the sequence of the synthetic promoters. The resulting plasmids were transformed into NEB5-alpha *F’ I*^*q*^ for their replication and storage. The new constructions were named pHC_DYO*DBD*-R, where *DBD* was substituted by the name of the regulator whose operator boxes have been introduced in the promoter (*Table S2*).

The collection of NEB5-alpha *F’ I*^*q*^ (pHC_DYO*DBD*-R) strains was used in a two pronged approach to validate and compare the functionality of the synthetic promoters driving the transcription of the GFP-encoding reporter gene. On the first set of experiments, we assessed individually the relative activity (Arbitrary fluorescence units / OD600) of every promoter compared to *Ptac*. This set of experiments determined that the synthetic promoters were working within a range of one order of magnitude from the most to the least active (*Figure S1*). Pre-cultures of the 15 strains were individually grown overnight in LB at 37°C. The next day they were used to inoculate fresh LB cultures in a 96-well plate (Flat bottom Polystyrene Black with clear bottom; Corning, Corning, NY) incubated at 37°C in a Synergy H4 Hybrid Multi-Mode Microplate Reader (Biotek, Winooski, VT), where OD600 and fluorescence in the emission range of GFP (excitation 485nm, emission 528 nm) were monitored along a time course of 12 to 24 hours. NEB5-alpha *F’ I*^*q*^ (pHC_DYOLacI-R) was included in every plate as the reference strain. Since the host strain expressed LacI these cultures were supplemented with 1 mM IPTG to prevent it from repressing the *Ptac* promoter driving GFP.

On the second set of experiments the 15 different NEB5-alpha *F’ I*^*q*^ (pHC_DYO*DBD*-R) strains were either grown separately in the same conditions described above or inoculated and grown together. Fresh 3-hour old cultures containing a mix of strains carrying the reporter plasmids were sorted in an Avalon Cell Sorter (PROPEL labs, Fort Collins, CO). Cells returning the strongest fluorescent emission signal for GFP (approximately 3-5% of the total) were recovered and plated on LB agarose plates containing carbenicillin. Individual clones were selected and the reporter promoters were identified by Sanger sequencing. The promoter abundance in the pre-sorting library was also characterized through Sanger sequencing of individual clones. The relative distribution of the promoters in the library before and after the selection process showed to be not statistically different once *Ptac* was removed from the data set, suggesting that the only preferentially enriched construction due to a stronger basal activity would be the unmodified *Ptac* promoter.

### Transformation of the pHC_DYO*DBD*-R collection of reporter plasmids into *E. coli* (Ch-END) and *E. coli* (Ch-OD)

Once the library of chimeric TFs was cloned into pCKTRBS/pCKTRBS-OD and the collection of reporter promoters was integrated in the pHC_DYO*DBD*-R vectors, it was necessary to obtain *E. coli* cells carrying one plasmid of each kind together. The screening method implemented in this work was based in a two-step FACS enrichment, with no selection marker pairing the DBD of a given chimera with its tailor-made reporter promoter. Both pCKTRBS/pCKTRBS-OD and pHC_DYO*DBD*-R carried antibiotic resistance markers, Cm^R^ and Ap^R^ respectively, but these genes were exclusively involved in the permanence of the plasmids in the host strain. As the copy-number of the pUC19 based pHC_DYO*DBD*-R vectors was higher than that of the pCK01 based pCKTRBS/pCKTRBS-OD, the former needed to be transformed into *E. coli* (Ch-END)/(Ch-OD) cells carrying the latter. The maximum possible efficiency was required for the transformation in order to increase the chances of generating a population of cells in which chimera and reporter were properly paired. Preliminary attempts electroporating ^81^ the reporter plasmids on the library of chimeras returned a very poor coverage (data not shown). The solution consisted in the generation of *E. coli* (Ch-END) and *E. coli* (Ch-OD) competent cells following the RbCl procedure ^69^. The efficiency of the newly competent cells was 10^8^-10^9^ cfu/μg. Three aliquots of competent *E. coli* (Ch-END)/(Ch-OD) cells were independently transformed with 100 ng of every pHC_DYO*DBD*-R plasmid resulting in an average 8.0 ± 2.9-fold coverage of the library per individual reporter plasmid. Transformed cells were pooled and selected in liquid adding chloramphenicol and carbenicillin to the culture media. After 7 hours of liquid selection the bacteria were preserved at -80°C. The two new libraries obtained after the recovery were identified as AYC Lib-Ch-END, short name for NEB5-alpha (pCKTRBS-*Chimera*, pHC_DYO*DBD*-R), and AYC Lib-Ch-OD, moniker for NEB5-alpha (pCKTRBS-*Chimera*-OD, pHC_DYO*DBD*-R), where *Chimera* represents any Nt-DBD-(LNK)-SBD-Ct.

### Characterization of relative abundance of chimeric TFs in bacterial populations

In test experiments where the relative abundance of a single chimeric TF compared to the rest of the library was required but a deep profile of the library was not needed, the method of choice was Sanger sequencing^82^of the pCKTRBS/pCKTRBS-OD vectors using the oligonucleotides pCKPolyF1 and pCKPolyR1 (*Table S3*). Pre and post-sorting samples were streaked on LB plates supplemented with the appropriate selection antibiotics and incubated at 37°C in order to obtain individual clones. A random set of clones of each condition was picked on a replica plate and once the colonies had grown the plate was submitted to Genewiz Inc. (Cambridge, MA) for the sequencing of the plasmids.

In some occasions it was needed to estimate the profile of chimeric TFs cloned into pCKTRBS/pCKTRBS-OD in a more thorough way. As a first step the plasmids present in the libraries were purified using commercially available kits (Qiagen, Hilden, Germany). These plasmid libraries served as template for the amplification of the joint region connecting the different domains that constitute the chimeric TF. For every chimera a region spanning from DBD-3’ to 5’-SBD was PCR amplified using the oligonucleotides shown in (*Table S3*). The Forward oligonucleotides hybridized at positions close to the 5’-end of the DBD-CORE and the reverse oligonucleotides hybridized upstream of the SP region of the SBD. The resulting PCR products contained enough information to identify DBD-CORE, DBD-END, LNK (if present), SBD-SP (if present) and SBD. In order to minimize PCR amplification bias based on amplicon size the reaction was monitored in real time as described above. Both the forward and reverse oligonucleotides contained flanking sequences enabling barcoding at both ends for pool paired-end sequencing performed in the MiSeq platform (Illumina, San Diego, CA). The resulting reads were analyzed using the DNA alignment program Bowtie ^83^. Subsequent handling of the data was performed using R (href="https://www.r-project.org/) in the RStudio environment (RStudio Inc., Boston, MA).

### FACS-based enrichment of benzoate-binding chimeric TFs

Fresh cultures of *E. coli* AYC Lib-Ch-END and AYC Lib-Ch-OD were started from overnight pre-cultures. Bacteria were grown in LB supplemented with chloramphenicol, carbenicillin and aTc. When the induction of the control chimera LacI-GGBP-OD was required the culture was supplemented with 0.4% glucose. In the screening for benzoate-binding TFs the cultures were supplemented with 1 mM benzoate. After growing for 4 hours, 2 ml of the cultures were precipitated (1 minute at 14000 rpm in a standard benchtop minifuge). Supernatants were removed and pellets resuspended in 1 ml PBS pH 7.2. Samples were diluted in PBS pH 7.2 at a density of 10^7^ cells/ml and sorted on a SH800Z Cell Sorter (Sony Biotechnology, San Jose, CA) using a 100 μM sterile sorting chip. The gating strategy (*Figure S4-A*) consisted of a population level round gate on a bivariate plot of forward scatter (FSC) area versus back scatter (BSC) area, followed by a rectangular gate containing various percentages of Enhanced Green Fluorescence Protein-Positive cells (EGFP^+^) on a bivariate plot of FSC area versus EGFP area. Excitation laser emission was 488 nm. Emission was collected with a 525/50 band pass (BP) filter. Optical layout below. (*Figure S4-B*) The sort mode was “Ultra Purity” and the event rate was approximately 5,000 per second. Cells were sorted into PBS pH 7.2 buffer.

### Cloning of the benzoate-responsive chimeric repressors ChTFBz01 and ChTFBz02

The ChTFBz01 (CbnR-ABE44898-OD) gene (*Supplementary Materials*) was ordered split on two synthetic dsDNA fragments (IDT, Coralville, IA). The first of them was flanked in its 5’-end by a 30-bp consensus RBS sequence included in pCKTRBS-OD (5′-CGGTACCCGGGTGACCTAAGGAGGTAAATA-3′) and PCR amplified with the primer pair RBS-Tail_F1 / cbnR_core-ABE44898-OD R1. The second fragment was flanked in its 3’-end by a 30-bp homology arm to the OD domain included in pCKTRBS-OD (5′-AAAAGAAAAACCACCCTGGCGCCCAATACG-3′) and PCR amplified with the primer pair cbnR_core-ABE44898-OD F1/END-OD-Tail_R1. A 30-bp homology region present in both the 3’-end of the first fragment and the 5′-end of the second fragment allowed for a correct cloning of the chimeric gene through a three-way Gibson assembly with a linear pCKTRBS-OD plasmid obtained by divergent PCR amplification (pCKTRBS RBSTail R1 / pCKTRBS-OD END-OD Tail F1). The assembled product was named pCKTRBS-CbnR-ABE44898-OD and transformed into chemically competent NEB5-alpha cells (NEB, Ipswich, MA). Once the chimeric construction was verified by Sanger sequencing the reporter plasmid for DBD-CbnR was electroporated into the validated chimera-bearing clone giving place to the NEB5-alpha (pCKTRBS-CbnR-ABE44898-OD, pHCDYOCbnR-R) strain. This strain was named AYC ChBzTF01.

The ChTFBz02 (LmrR-BzdB1_nSP) gene (*Supplementary Materials*) was ordered as a single synthetic dsDNA fragment (IDT, Coralville, IA) flanked in its 5′-end by the 30-bp consensus RBS sequence mentioned above and in its 3′-end by a 30-bp sequence present on pCKTRBS polylinker (5′-GATCCTCTAGAGTGGACCTGCAGGCATGCA -3′). The fragment was PCR amplified using the primer pair RBS-Tail_F1 / END-Tail_R1 and Gibson assembled into a linear pCKTRBS vector obtained by divergent PCR using the primer pair pCKTRBS RBS-Tail R1 / pCKTRBS END-Tail F1. The assembled product was named pCKTRBS-LmrR-BzdB1_nSP and transformed into chemically competent NEB5-alpha cells (NEB, Ipswich, MA). Once the chimeric construction was verified by Sanger sequencing the reporter plasmid for DBD-LmrR was electroporated into the validated chimera-bearing clone giving place to the NEB5-alpha (pCKTRBS-LmrR-BzdB1_nSP, pHC_DYOLmrR-R). This strain was named AYC ChBzTF02.

### *In vivo* assay of ChTFBz01 and ChTFBz02 activity

AYC ChTFBz01 and AYC ChTFBz02 strains were used to assay the *in vivo* functionality of the new chimeric repressors. Pre-cultures of the two strains were individually grown overnight in LB medium. The next day they were used to inoculate fresh M63 minimal medium cultures in a 96-well plate (Flat bottom Polystyrene Black with clear bottom; Corning, Corning, NY) that was in turn incubated at 37°C in a Synergy H4 Hybrid Multi-Mode Microplate Reader (Biotek, Winooski, VT), where OD600 and fluorescence in the emission range of GFP were monitored along a time course for a minimum of 8 hours.

## ABBREVIATION LIST

3D: Three-dimensional
Apm^R^: Resistance to ampicillin/carbenicillin
aTc: Anhydrotetracycline
Cm^R^: Resistance to chloramphenicol
CRISPR: Clustered Regularly Interspaced Short Palindromic Repeats
Ct: C-terminal
DBD: DNA-Binding Domain
dCas9: Defective Cas9 protein, able to bind DNA and RNA but lacking nuclease activity
eLCR: enhanced LCR
FACS: Fluorescence-Associated Cell Sorting
GGBP: glucose-binding protein (*E. coli*)
Gm^R^: Resistance to gentamycin
IPTG: Isopropyl β-D-1-thiogalactopyranoside
Km^R^: Resistance to kanamycin
LCR: Ligase Chain Reaction
LNK: linker
nSP: SBD that does Not contain its Signal Peptide
Nt: n-terminal
PBP: Periplasmic Binding Protein
PBS: Phosphate Buffered Saline
RBS: Ribosome Binding Site
SBD: Substrate-Binding Domain
OD: Oligomerization Domain
PID: Percentage of Identity
SP: Signal Peptide
TALENs: Transcription activator-like effector nuclease
TF: Transcription factor

## ACKNOWLEDGEMENTS

This material is based upon work supported by the U.S. Department of Energy (DOE), Office of Science, Biological and Environmental Research Program under Award Number DE-FG02-02ER63445 (PI G.M.C.). We want to acknowledge H.H. Lee, R. Kalhor, N. Ostrov and A.H. NG (Harvard Medical School) as well as I.F. Escapa (Forsyth Institute) for their help discussing and reviewing this work. We thank Dr. F.J. Carrillo-Salinas (Tufts University School of Medicine) for the graphic design of *Figure 2*, D.J. Wiegand and B. Turczyk (Wyss Institute) for the CustomArray DNA synthesis, J.K. Moore and S. Terrizzi (Harvard medical School) for their assistance with FACS, N. Eroshenko (Harvard Medical School) for his oligonucleotide collection and B.L. Wanner (Harvard Medical School) for sharing his vectors. J.F. Juarez thanks J. Aach (Harvard Medical School) and V. Raman (University of Wisconsin-Madison) for the seminal discussions that originated this paper and E. Diaz, G. Durante-Rodriguez and M. Carmona (CIB-CSIC) for their lessons on chimeric genes.

